# Investigating the dynamics of proviral silencing in polyclonal HIV-1 infected Jurkat cell populations

**DOI:** 10.64898/2026.02.26.708239

**Authors:** Shelby Clark, Edmond Atindaana, Kamya Gopal, Jeffrey M Kidd, Alice Telesnitsky

**Affiliations:** Department of Microbiology & Immunology, University of Michigan Medical School, Ann Arbor, Michigan, USA; Cellular and Molecular Biology Program, University of Michigan Medical School, Ann Arbor, Michigan, USA; Department of Human Genetics, University of Michigan Medical School, Ann Arbor, Michigan, USA

## Abstract

Previous work demonstrated that individual HIV-1 provirus-containing Jurkat cell clones maintain stable bimodal expression patterns throughout 8 to 9 days of culture. However, within polyclonal infected cell pools, the overall proportion of cells displaying HIV-1 expression declines with time. Here, we examined the processes underlying population silencing in polyclonal pools comprised of hundreds of individual barcoded proviral clones throughout culturing periods of 22 and 90 days. Monitoring HIV-1 LTR activity via a GFP reporter confirmed that initially most clones exhibited bimodal expression, with mixtures of transcriptionally active and inactive cells. Over time, however, the overall fraction of LTR-active cells declined substantially. High-throughput clonal tracking showed that this decline was not driven by uniform transcriptional silencing. Instead, population silencing resulted from a combination of mechanisms: selective expansion of clones with low LTR activity, reductions in expression within certain clones, and long-term maintenance of stable bimodal expression in others. Notably, over half of clones retained stable bimodal expression patterns after 22 days, and approximately 17% of the clones retained stable bimodal expression even after 90 days in an independent second experiment, despite ongoing phenotypic switching at the single-cell level. These findings demonstrate that population-level HIV-1 silencing emerges from heterogeneous, clone-specific behaviors rather than uniform transcriptional repression. These observations confirm stability for a large subset of clones and provide insight into how integrants with diverse replication and expression properties can act together during HIV-1 polyclonal population silencing.

## Introduction

The half-lives of most cells productively infected with HIV-1 are only about two days^1^. Nonetheless, because HIV-1 integrates its genome into the host cell’s DNA, the virus can persist for decades in cells and their daughters when cells survive initial infection. Antiretroviral therapy (ART), the current standard of care for HIV-1 infection, prevents cell-free virus dissemination, effectively suppressing viremia to below detectable limits. However, when ART is discontinued or interrupted, viral rebound occurs and rekindles infection, therefore requiring lifelong treatment for people living with HIV ^2, 3^. The major source of viral rebound is believed to be a residual HIV-1 reservoir that persists during therapy and consists of a small fraction of infected cells harboring transcriptionally silent but replication-competent proviruses^4^. Despite decades of research, the mechanism governing reservoir formation, transcriptional regulation, and long-term persistence remain poorly understood.

Multiple factors have been suggested to contribute to the establishment and maintenance of the reservoir. Transition of infected CD4+ T cells into a resting memory state, which results in reduced availability of key transcription factors such as NFkB, SP-1, and p-TEFB, creates a cellular environment that favors transcriptional repression^5, 6^. Deficiencies in viral regulatory proteins, particularly the HIV transcriptional transactivator protein Tat, can similarly promote entry into a silent state ^7, 8^. Epigenetic modification of the local chromatin environment, including histone acetylation and methylation, further influences whether proviruses remain active or are transcriptionally silenced. Integration site features, including genomic location, provirus orientation within genes, and proximity to regulatory elements, have also been implicated in determining whether a provirus becomes long-lived ^4, 9-11^.

Importantly, studies of individuals on suppressive ART have revealed that the reservoir is polyclonal but dominated by groups of infected cells carrying identical proviral integration sites, implying that clonal expansion of a small fraction of proviruses contributes substantially to reservoir stability. This clonal expansion is thought to arise through normal T-cell homeostatic proliferation, antigen-driven responses, or cell intrinsic differences that enhance the fitness of particular proviral integrants ^12, 13^.

Experimental studies have demonstrated that HIV-1 infection generates a heterogenous population of provirus-containing cells with remarkably variable HIV-1 expression ^14^. Single proviruses isolated from people living with HIV-1 also show dramatic differences in virion production, with levels of virus production differing among integrants by up to 100,000-fold^15^. Even within clonal cell lines, individual proviral clones display bimodal expression patterns in which dividing cell population members fluctuate between transcriptionally active and inactive states. These heterogenous expression patterns are influenced by stochastic noise in gene regulation, driven in part by Tat-mediated positive feedback loops and host transcriptional variability ^16^. Previous work tracking individual HIV-1 infected clones has demonstrated that despite dynamic oscillations in LTR activity, many proviruses exhibit heritable, clone-specific bimodal expression patterns over a two-week period ^17^. These observations suggest that HIV-1 gene expression is governed by a combination of stochastic and deterministic factors acting in a clone intrinsic manner. However, notwithstanding the above mentioned work demonstrating the maintenance of HIV-1 expression throughout 8 to 9 days of Jurkat cell culture, proviral silencing is a widely recognized phenomenon and polyclonal pools of infected Jurkat cells silence gradually over prolonged passaging ^18^.

In the current study, the dynamics of proviral silencing of hundreds of individual clones within polyclonal HIV-1 infected Jurkat cell pools were monitored. Using a high throughput barcoding and sequencing approach, the contributions of clonal expansion as well as changes in bimodal expression patterns were investigated over an extended duration of passaging. The findings indicated that the initial increase in population silencing was largely driven by the clonal expansion of a few low-LTR active clones rather than transcriptional silencing of individual proviruses. However, at later time points, cell loss of LTR-driven fluorescence reflected a more complex combination of factors, including clonal expansion of low-LTR active clones, loss of high-LTR active clones, transcriptional silencing of a subset of individual proviruses, and maintenance of bimodal proportions by some clones. Together, these results illustrate how clones with diverse expression and replication properties interact in cell culture to generate long-term shifts in HIV-1 polyclonal populations.

## Results

### Monitoring population silencing in barcoded polyclonal HIV-1 infected Jurkat T cell pools

To investigate the dynamics of population silencing within a polyclonal HIV-1 infected pool, Jurkat T cells were infected with an NL4-3 based vector encoding all HIV-1 proteins except for Env, Vpr and Nef (Fig. 1A). Due to a deletion in the *env* region, the virus is replication deficient and undergoes only a single cycle of replication. A green fluorescent protein (eGFP) reporter driven by the HIV-1 LTR was inserted into the *nef* open reading frame to monitor provirus transcriptional activity. The presence of a puromycin resistant gene driven by the EF1α promoter enabled selection of infected cells independent of LTR activity. A 20-bp randomized sequence tag or ‘barcode’ was incorporated into the U3 region of the 3’ LTR. Upon reverse transcription this tag was duplicated to both LTRs and fixed at the integration site, thereby serving as a unique clonal identifier.

**Figure 1:**
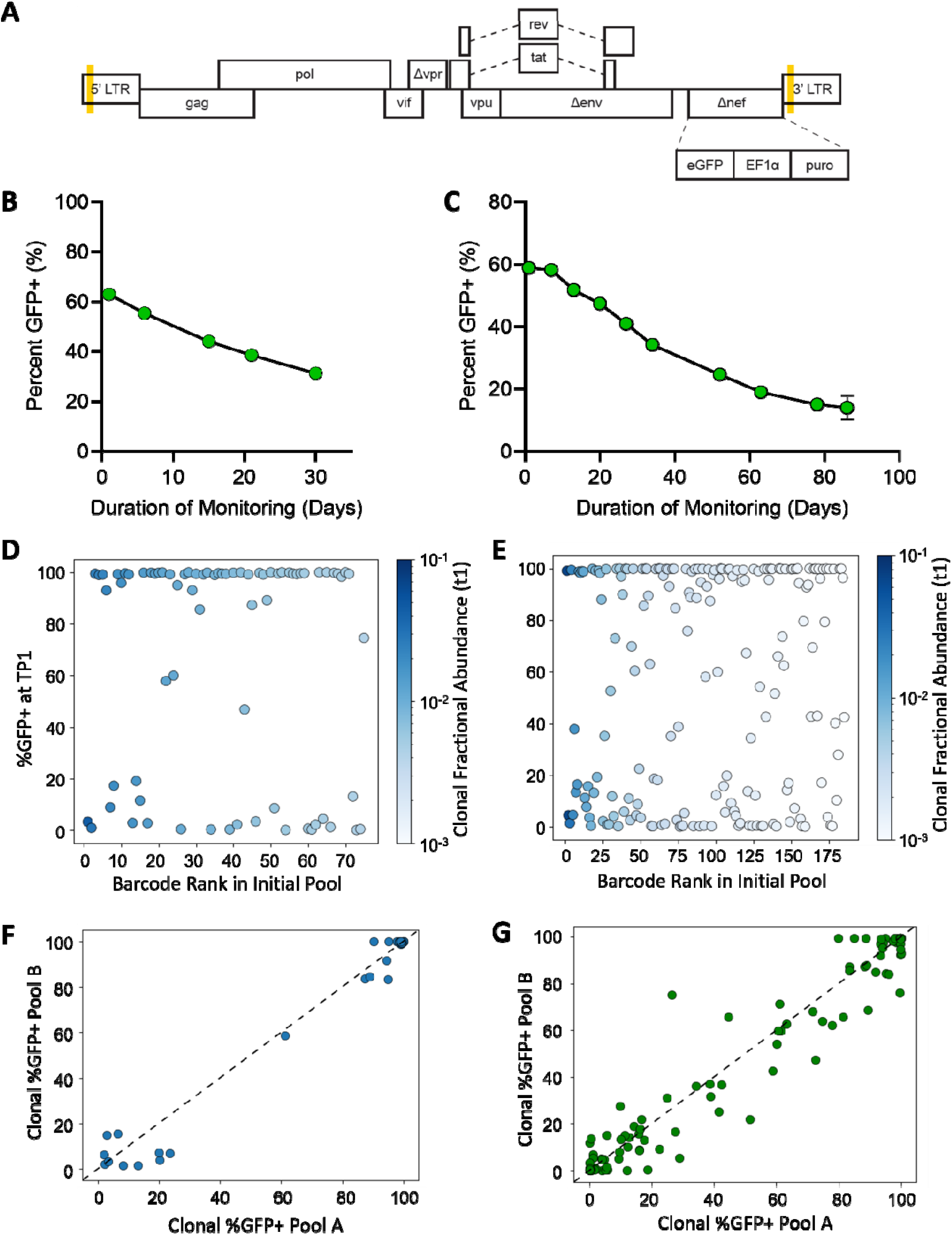
Monitoring population silencing in polyclonal HIV-1 infected Jurkat T cell pools. (A) Schematic of HIV-1 vector used to produce barcoded virus. A 20-bp randomized barcode was placed in the U3 of the 3’ LTR and duplicated in both LTRs following reverse transcription (indicated by yellow shading). Enhanced green fluorescent protein (eGFP) plus eF1α promoter driving the constitutive expression of the puromycin resistance gene was inserted within the *nef* open reading frame. (B, C) Representative line graph depicting %GFP+ of unsorted polyclonal barcoded HIV-1 infected Jurkat T cell population at indicated day post start of monitoring as assessed by flow cytometry for (B) Pool 22d and (C) Pool 90d. (D, E) Clonal %GFP+ in (D) Pool 22d and (E) Pool 90d. Clonal %GFP+ calculated by dividing read frequency of each barcode in GFP+ sorted subpool by the sum of the abundance in GFP+ and GFP− subpools after weighting to proportions of total cells in GFP+ and GFP− sorted pool as determined by flow cytometry. Barcode rank refers to where that clone ranked when barcodes were listed in order of total cell abundance at initial timepoint. Clones are colored based on fractional abundance at t1 as indicated by the color bar. (F, G) Comparison of %GFP+ determined for each barcode in Pool A and Pool B of (F) Pool 22d (Spearman ρ = 0.718054, p= 1.93 × 10^−08^, n = 51) and (G) Pool 90d 22d (Spearman ρ = 0.911933, p= 3.48 × 10^−57^, n = 145).

Previous work using this barcoded HIV-1 system showed that individual clones maintained stable, clone-specific mixtures of GFP− and GFP+ cells despite passaged cells alternating between LTR-active and LTR-inactive states ^17, 19^. These equilibrium proportions were heritable and persisted over time. Continued culture revealed a gradual decrease in the proportion of infected cells that displayed LTR activity, suggesting that the system can be used to monitor dynamics of population silencing within a polyclonal HIV-1 infected pool.

To quantify silencing dynamics, two independent polyclonal pools were generated here, each by infecting 5 × 10^6^ Jurkat T cells at a low MOI (0.0005) to ensure single integration events. One polyclonal pool was maintained for 22 days (Pool 22d) and the other was maintained for 90 days (Pool 90d). Following puromycin selection, both pools were maintained without drug and divided into two replicate cultures 14 days after initial infection, with the replicates subsequently maintained separately (replicates A and B). LTR activity of the populations, measured as a percentage of GFP positive cells (%GFP+), was measured periodically from the initial day of monitoring (t1) to the end of passaging by flow cytometry (Fig. 1B, 1C). Initially, replicates 22d-A and -B both contained approximately 64% GFP+ cells. By day 22 (t22), the GFP+ fractions of these pools had decreased to approximately 38% (Fig 1B). Replicates 90d-A and -B contained approximately 60% GFP+ cells at the beginning of monitoring (t1) and decreased to approximately 18% GFP+ cells after 90 days (t90) (Fig. 1C). Pool 22d at t1 and t22, and Pool 90d at t1, t22, and t90, were sorted into GFP+ and GFP− fractions. At each of these timepoints, cell genomic DNA was isolated from the sorted subpools and from an unsorted aliquot of each cell population.

High-throughput sequencing of DNA from 2 × 10^6^ cells at t1 identified 370 unique barcodes for Pool 22d and 542 unique provirus barcodes for Pool 90d (data available upon request). For downstream analysis, only the most abundant clones—those with a fractional abundance greater than 0.001 of the total pool—were included to ensure robust and biologically meaningful comparisons. Bimodal expression patterns for these well-represented clones (75 clones for Pool 22d and 185 clones for Pool 90d) were calculated by dividing the read frequency of each barcode in the GFP+ sorted sub pool by the sum of the abundance in both the GFP+ and GFP− subpools. Values, designated as clonal %GFP+, were weighted to reflect the proportions of total cells in the GFP+ and GFP− sub pools as determined by flow cytometry.

Both Pool 22d and Pool 90d displayed broad distributions of clonal %GFP+ and included clones ranging from >99% to <1% GFP+ (Fig. 1D, 1E). Note that the rank order of clones reflects their fractional abundance within the polyclonal populations at t1. Consistent with previous work using this system, most clones displayed either high (>80%) or low (<20%) proportions of GFP+ cells, with relatively few clones displaying intermediate values, and the proportion of clones displaying low %GFP+ values was similar among abundant and rarer clones, suggesting HIV-1 gene expression levels were not responsible for clonal differences in bimodal proportions at t1 ^17^.

To determine whether these %GFP+ values reflected true clone-specific expression patterns rather than sampling variability, clonal %GFP+ were calculated independently for replicate pools A and B and plotted against one another (Fig. 1F, 1G). Clonal %GFP+ were highly concordant between replicates (Pool 22d: Spearman ρ = 0.718, p = 1.93 × 10^−08^, n = 51; Pool 90d: Spearman ρ = 0.912, p = 3.48 × 10^−57^, n = 145), indicating that the observed expression patterns were not attributable to sampling noise. These results confirmed that the barcode sequencing approach provides a robust method for tracking individual proviruses within a polyclonal population and quantifying their contributions to population-level expression. Based on these findings and sustained similar values for A and B datasets for each timepoint, data for replicates A and B were combined in subsequent analyses presented below.

### Stable bimodal expression patterns observed in approximately 52% of Pool 22d clones after 22 days of passage

To determine whether population-level silencing arose from clone-intrinsic transcriptional changes, shifts in clonal representation, or a combination of both, the dynamics of bimodal expression for each proviral clone within the polyclonal pools were monitored over time. To achieve this, the log change in clonal %GFP+ between t1 and t22 for Pool 22d and the changes for Pool 90d at t1, t22, and t90 were quantified (Fig. 2A, 2B, 4A, 4B). This analysis revealed that despite an overall decrease from 64% to 38% in GFP+ expression at the population level, approximately 52% of the clones that remained detectable in Pool 22d at t22 maintained stable bimodal expression patterns, defined as a log change in clonal %GFP+ within ±0.05. Larger shifts in %GFP+ were observed with the remaining clones: in most cases these were reduced %GFP+ levels (Fig. 2B). As another way of examining these trends, the %GFP+ values for each clone were compiled at t1 and at t22 and mean values for each clone at each timepoint were determined (Fig. 2C). This comparison revealed only a modest decrease in the mean %GFP+ when expression was considered at the clonal level, indicating that the population-level decline in LTR-driven fluorescence could not be explained by clone-intrinsic transcriptional silencing alone.

**Figure 2:**
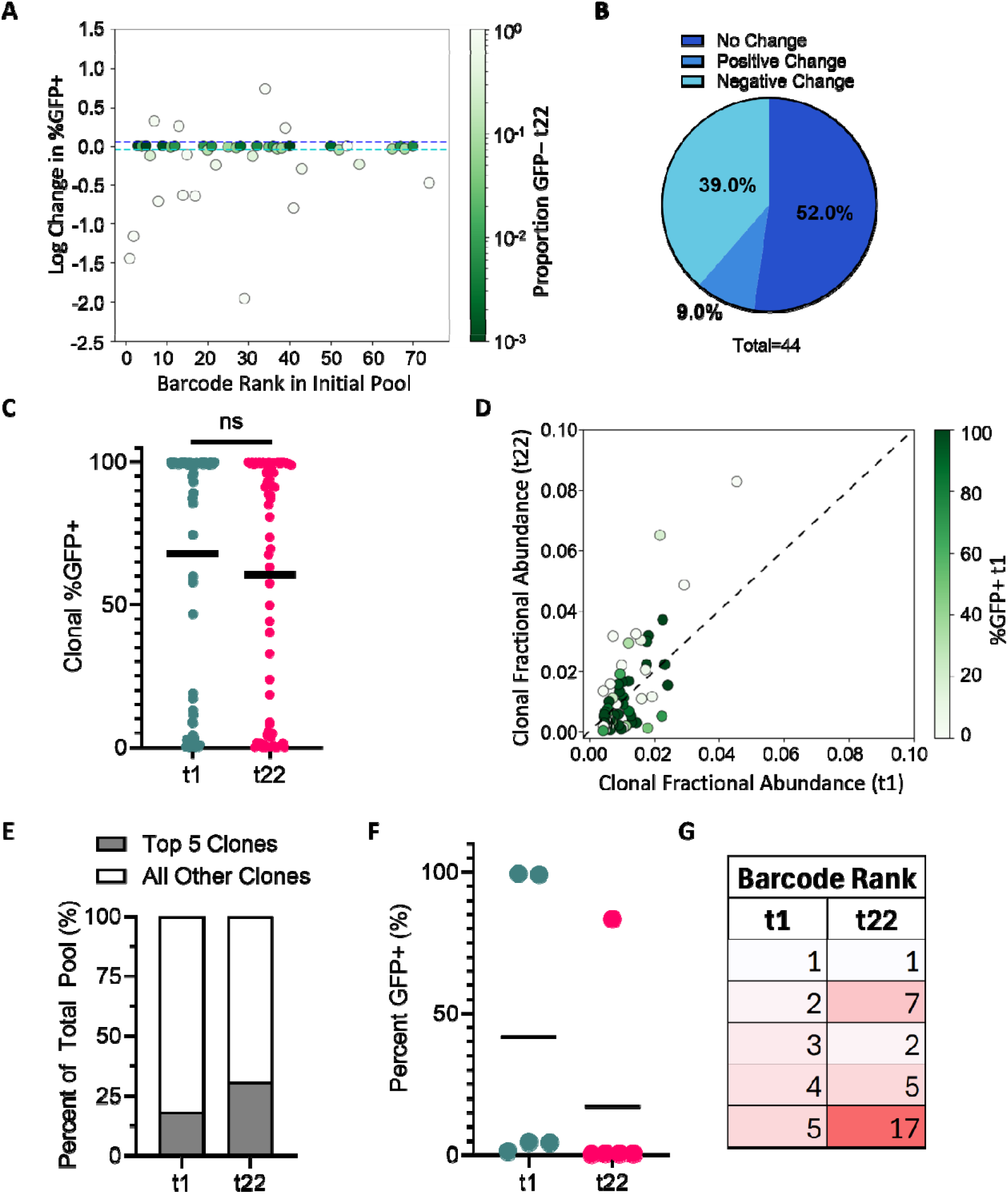
Stable bimodal expression patterns observed for approximately 52% of the proviral clones over 22 days of passaging. (A) Change in clonal %GFP+ in Pool 22d. Y-axis indicates log change in clonal %GFP+ from t1 to t22. Clones are colored based on proportion of GFP− at t22 (0.001-1.0). Colored dash lines indicate limits for determining positive (dark blue) and negative (light blue) change. (B) Percentage of clones exhibiting no change (log change between ±0.05), positive change (>0.05) or negative change (<-0.05) in %GFP+. (C) Scatter dot plot depicting clonal %GFP+ at t1 and t22. Mean clonal %GFP+ indicated by horizontal bar. Differences in t1 and t22 clonal %GFP+ were compared using a Mann-Whitney U two-sided test, (ns, p = 0.1006). (D) Clonal fractional abundance at t1 versus t22. Clones colored based on clonal %GFP+ at t1. (E) Percent composition of the five most abundant clones versus all other clones of the polyclonal pool at t1 and t22. (F) %GFP+ for the five most abundant clones at t1 and t22. Horizontal line indicates calculated mean. (G) Barcode rank of the five most abundant clones at t1 and t22. Box shading reflects clone rank order at t1.

This conundrum was partially resolved by a comparison of clonal fractional abundances in Pool 22d at t1 and t22, which showed that a small subset of clones expanded disproportionately over time (Fig. 2D). This analysis revealed that at t1, approximately 18% of the cells in Pool 22d was comprised of the five most abundant clones, with the relative proportion of the top five clones increasing to approximately 30% of the polyclonal population by t22 (Fig. 2E). Notably, the five most abundant clones at t22 were not the same clones that dominated at t1. Instead, several clones that were initially lower ranked (notably clones #7 and #17) expanded more rapidly, displacing two of the top five clones present at t1 (Fig. 2G). These expanded clones were predominantly low–LTR-active clones, demonstrating that their outgrowth contributed substantially to the population-level decline in fluorescence (Fig. 2F).

The stability of bimodal expression properties for individual cell lineages in Pool 22d was assessed by maintaining the sorted GFP− and GFP+ subpools separately for 22 days, after which each subpool was resorted into GFP+ and GFP− populations (Fig. 3A). Analysis of clonal %GFP+ within each resorted subfraction demonstrated that not all descendants of GFP− cells remained GFP−, and not all descendants of GFP+ cells remained GFP+, indicating ongoing oscillation between LTR-active and LTR-inactive states during cell passage (Figure 3B, 3C). Despite this phenotypic switching, computational reconstruction of the unfractionated pool, generated by combining weighted data from the fractions resorted after 22 days, strongly correlated with the clonal %GFP+ values measured in the original unsorted pool (Spearman ρ = 0.826, p = 7.64 × 10^−20^, n = 75; Fig. 3D). These results are consistent with those we previously reported for cells that were passaged for 8 to 9 days and demonstrate stable inheritance of clone-specific expression patterns for many clones, and the appearance of phenotypic drifting for most other clones, despite dynamic single-cell transitions between active and inactive states over 22 days ^17^.

**Figure 3:**
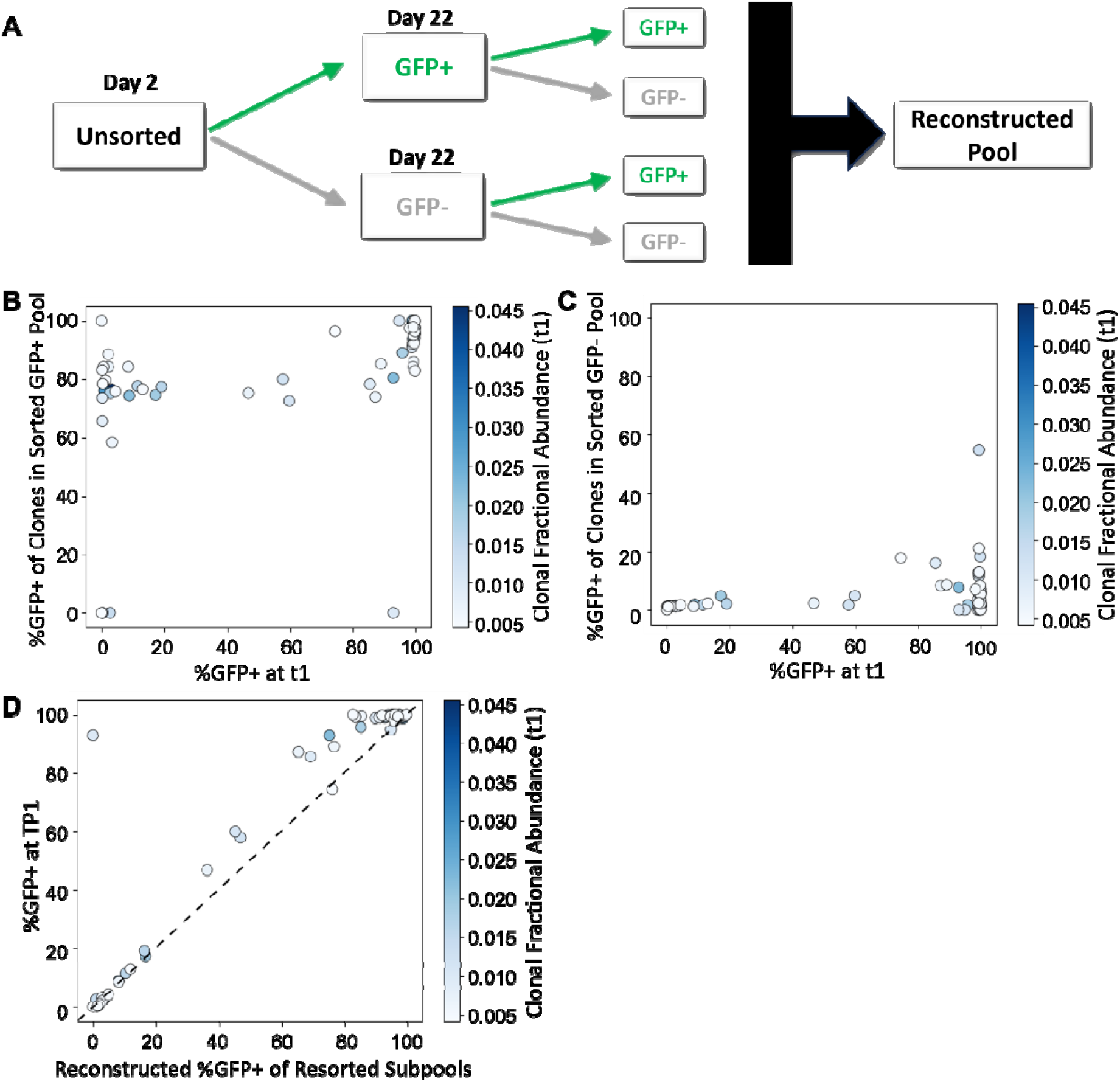
Bimodal expression patterns remain stable over 22 days despite observed phenotypic switching of cells between LTR active and LTR inactive. (A) Schematic of passaging and sorting of polyclonal pools, with initial sorted pools followed by the resorted subpools. Reconstructed pool calculated by weighting GFP percentage of resorted subpools against initial unsorted pool. (B, C) Calculated %GFP+ of Pool 22d clones following resorting of (B) the GFP+ sorted sub pool and (C) the GFP− sorted sub pool compared to clonal %GFP+ in initial unsorted pool. (D) Reconstructed clonal %GFP+ after resorting the GFP+ and GFP− subpools and weighting to initial pool, compared to clonal %GFP+ in initial unsorted pool. Clones are colored based on fractional abundance at t1 (Spearman ρ = 0.826, p = 7.64 × 10^−20^, n = 75).

### Stable bimodal expression patterns maintained in approximately 17% of Pool 90d clones after 90 days

We then moved on to analyze clone dynamics in an independently derived Jurkat cell library, called Pool 90d. 90d transitioning from approximately 59% GFP+ at t1 to approximately 47% GFP+ at t22 and approximately 64% of clones that maintained fractional abundance above 0.001 maintained stable bimodal proportions (Fig. 1C, 4H, and not shown). At t90, approximately 17% of detectable clones exhibited stable bimodal expression patterns, in that their %GFP+ proportions remained unchanged throughout the three-month experiment (Fig. 4A, 4B). However, the mean of the %GFP+ values for all clones decreased substantially from t1 to t90 (Fig. 4C). This decline reflected both the loss of highly active clones (“dropped” clones) and clone-intrinsic silencing among previously active clones. By t90, approximately 78% of t1 clones were no longer detected or had fallen below the set threshold for analysis (fractional abundance less than 0.001); these dropped clones were predominantly high-LTR active (Fig. 4C).

**Figure 4:**
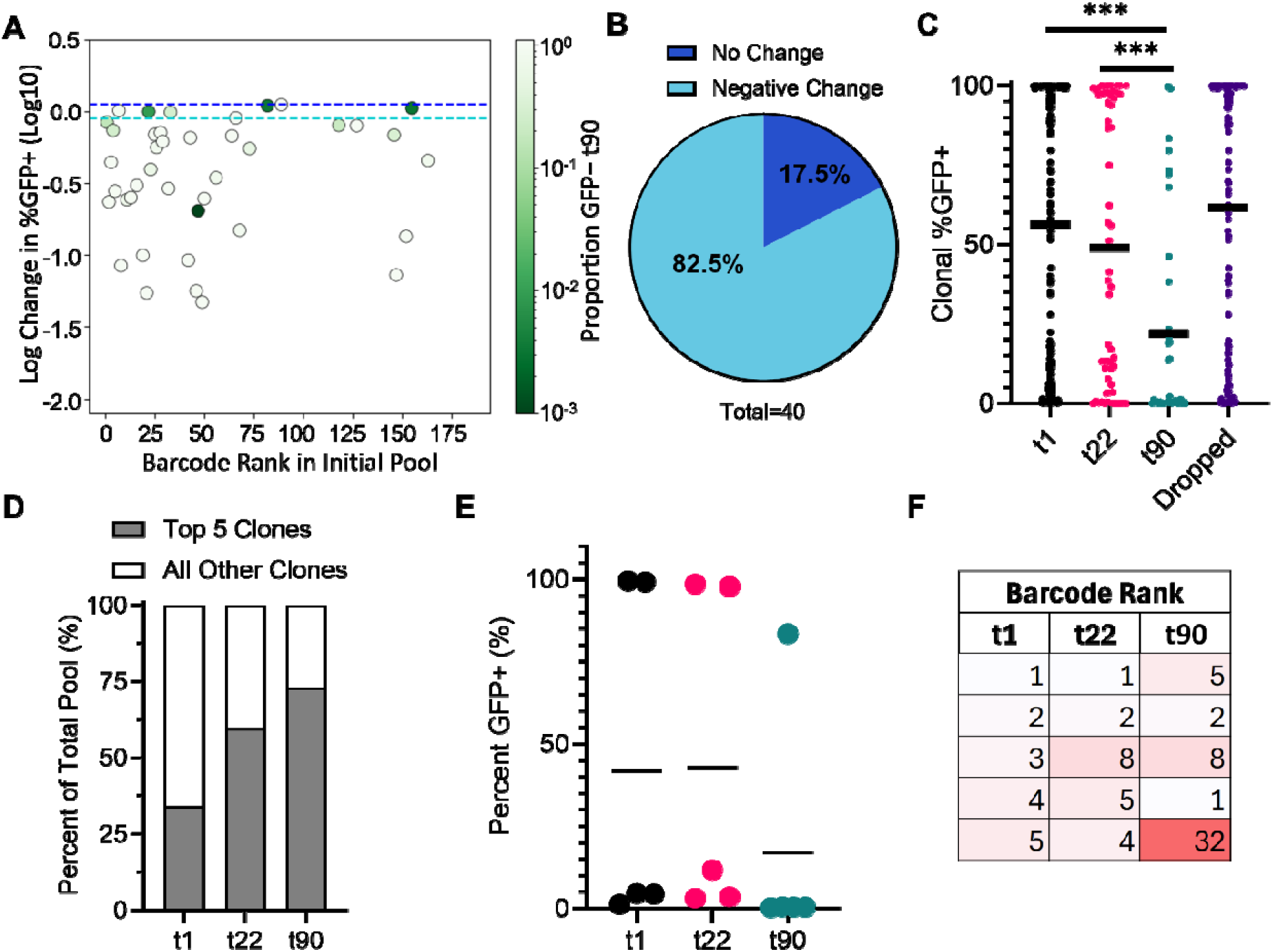
Stable bimodal expression patterns maintained in approximately 17% of clones after 90 days. (A) Change in clonal %GFP+ in Pool 90d. Y-axis indicates log change in clonal %GFP+ from t1 to t90. Clones are colored based on proportion of GFP− at t90 (0.001-1.0). Colored dash lines indicate limits for determining positive (dark blue) and negative (light blue) change. (B) Percentage of clones exhibiting no change (log change between ±0.05) or negative change (<-0.05) in %GFP+. (C) Scatter dot plot depicting clonal %GFP+ at t1, t22, t90 and dropped clones. Mean clonal %GFP+ indicated by horizontal bar. Differences in mean clonal %GFP+ were compared using a Mann-Whitney U two sided test, (t1 vs t22: ns, p= 0.1455, t1 vs t90: ***, p = <0.0001, t22 vs t90: ***, p= 0.0005). (D) Percent composition of the five most abundant clones versus all other clones of the polyclonal pool at t1, t22, and t90. (E) %GFP+ for the five most abundant clones at t1, t22, and t90. Horizontal line indicates calculated mean. (F) Barcode rank of the five most abundant clones at t1 t22, and t90.

Clonal fractional abundances again shifted toward a small group of expanding proviruses: the five most abun Pool 90d differed at t22 from t1 and at t90 from t22, with lower-ranked clones expanding more rapidly than early dominant clones (Fig. 4F). For most but not all silencing clones, t22 %GFP+ values were intermediate to those observed at t1 and t90. But when the dynamics of clone-intrinsic silencing was examined over the three tested timepoints, it was observed that not all clones silenced at the same rate (Fig. 5A, 5B). Together, these results demonstrate that population-level silencing properties were similar in two independently-derived pools, and that this silencing was driven primarily by the preferential expansion of a small number of low–LTR-active clones and, over extended passaging, by the additional effects of clonal loss and successive silencing of formerly active clones. However, the population also maintained a substantial subset of proviruses that retained stable, clone-specific bimodal expression patterns over time.

**Figure 5:**
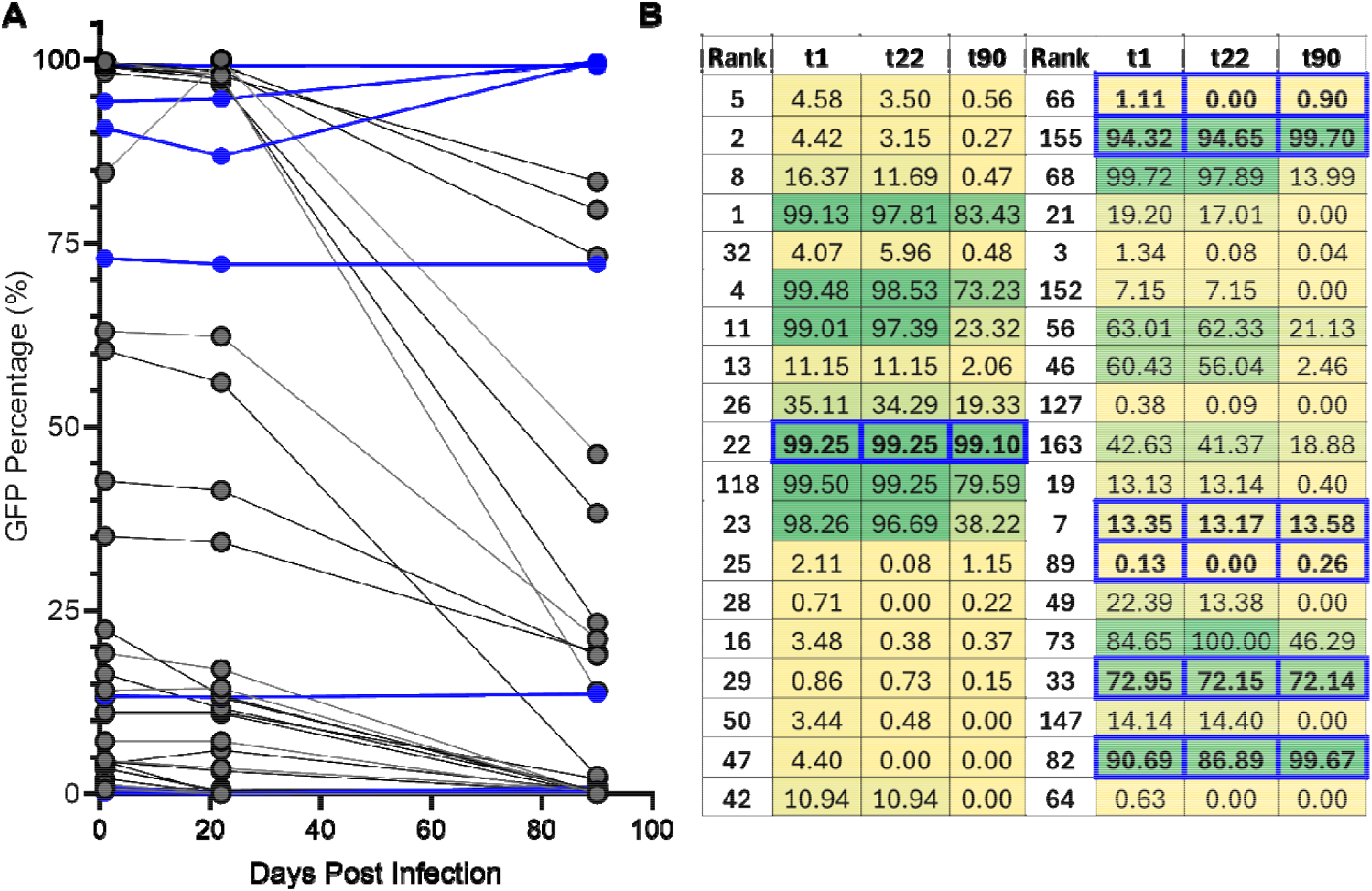
Clone-intrinsic silencing occurs at variable rates. (A) GFP+ percentage at t1, t22 and t90 of the 40 most abundant clones still detected by t90. Samples in blue indicate clones that maintained bimodal proportions throughout 90 days (log change in %GFP between +0.05 and −0.05). (B) Table depicting clonal %GFP at each timepoint of the 40 most abundant clones still detected by t90. Provided clone numbers are rank order at t1. The ordering in this panel reflects clone abundance at t90. Box shading reflects clonal proportions of GFP+ cells. Blue outlines indicate clones that maintained bimodal proportions throughout 90 days.

## Discussion

This study characterized the long-term dynamics of proviral expression and clonal expansion within polyclonal HIV-1–infected Jurkat T cell populations. Using a high-throughput barcoding approach, hundreds of individual proviral clones were tracked over 22- and 90-day culture periods and both LTR-driven GFP expression and clonal fractional abundance were quantified. Our findings demonstrate that population-level silencing arose from a combination of clone-intrinsic transcriptional behavior and selective clonal proliferation.

Consistent with previous reports of heritable, clone-specific LTR activity^17^, a substantial subset of proviruses maintained stable proportions of transcriptionally active and inactive cells over extended culture periods (Fig. 2, 3, 4). Approximately 52% of clones retained stable bimodal expression after 22 days, and 17% remained stable after 90 days. By resorting separately passaged GFP+ and GFP− subpools, we observed that although clonal %GFP+ proportions remaining stable in a subset of proviruses, daughter cells did not uniformly adopt the parental phenotype. Instead, cells alternated between GFP+ and GFP− states, consistent with previously described phenotypic bifurcation and stochastic fluctuations in HIV transcription (Fig. 3) ^16, 17^. Yet when clonal %GFP+ values from the reconstructed population at t22 were compared to the original unsorted values at t1, most clones remained highly concordant. This observation indicates that despite the stochastic variability at the single cell level, clone-specific expression equilibrium can be preserved over three weeks, and subsequent timepoints demonstrated that stable bimodal expression patterns were maintained for some clones throughout months of cell passage. These results reinforce the concept that HIV-1 gene expression is governed by a combination of stochastic and deterministic factors.

Future studies investigating the interplay between stochastic and deterministic inputs involved in each clone adopting a unique bimodal expression pattern are needed for parameters of both host and viral origin. We hypothesize that each clone adopts a unique expression pattern through multiple inputs, both stochastic and deterministic, that combine in clone-intrinsic ways to skew the probability that a given cell will reach the conditions needed for LTR expression. Previous work has explored host contributions to proviral expression levels that are related to integration site features including proviral orientation, as well as proximity to repressive or activating chromatin marks ^17, 20-24^. A significant difference in % LTR-active cells was observed by proviral orientation, but modest to negligible differences were noted for proximity to chromatin marks, suggesting that proximity to mapped chromatin features alone does not explain expression variability.

Although some individual clones maintained stable bimodal expression patterns, population-level %GFP+ declined substantially over time (Fig. 1). Our analysis revealed that this decline was largely attributable in early stages of passaging to selective expansion of low-LTR-active clones, rather than widespread transcriptional silencing of active clones. In both the 22- and 90-day cultures, the most abundant clones accounted for a disproportionately large fraction of the population at the end of cell passaging, and these clones were predominantly low–LTR-active (Fig. 2, Fig. 4). Over extended culture (90 days), additional factors contributed to population silencing. Some previously active clones were lost entirely (“dropped”), and another subset of initially active clones displayed reduced proportions of LTR active cells over time. When clonal silencing dynamics were examined, some clones maintained stable bimodal expression patterns, whereas others exhibited gradual declines with successive passages or remained stable initially before undergoing rapid silencing at later timepoints (Fig. 3). Note, however, that these observations are limited by the number of timepoints analyzed in the current study, as well as the modest read depth achieved with the sequencing approaches used, and more comprehensive longitudinal studies would be required to fully resolve the dynamics of clonal silencing. Together, these findings indicate that population-level declines in Jurkat provirus LTR activity over long durations were the result of a complex interplay of clonal expansion, clonal attrition, and intrinsic transcriptional silencing, rather than a single dominant mechanism in our system. To some extent, this mirrors in vivo observations that show the reservoir is dynamic, with certain clones expanding, contracting, or becoming latent over time ^13, 25^.

It is tempting to make inferences from this in vitro study to clinical observations. For example, disproportionate clonal expansion was observed in this Jurkat cell system, and clonal expansion is a major feature of the reservoir in patients on suppressive ART, where only a small number of infected T-cell clones dominate through selective proliferation^12^. However, our study was performed in Jurkat T cells, which are known to exhibit genomic instability and lack the complexity of *in vivo* reservoirs, including T cell developmental stage, interactions with the immune system, and tissue-specific microenvironments. Notably, approaches similar to those used here are common in cell-based models for HIV-1 latency, and the clone-specific differences and dynamics observed here may help explain some of the variability intrinsic to those systems ^26, 27^.

Together, the findings here highlight that heterogeneous proviral populations are governed by layered regulatory processes operating at both single-cell and clonal scales. Although the experimental design of removing *env, vpr*, and *nef* was intended to minimize the presence of cytotoxic HIV-1 gene products, some of our observations may reflect residual effects of HIV-1 gene expression. Notably, early population silencing in this system was driven primarily by differential proliferative fitness, favoring expansion of low–LTR-active clones, while long-term silencing emerged through the combined effects of clonal attrition and transcriptional shutdown of previously active proviruses. These insights underscore the need for future studies to dissect the viral and host determinants that set and stabilize HIV-1 expression patterns and define clonal fitness.

## Methods

### Cell Lines and Cell Culture

Human embryonic kidney HEK293T cells and Jurkat T cells were obtained from the American Type Culture Collection (ATCC, Rockville, MD, USA; CRL-3216 and TIB-152, respectively). HEK293T cells were cultured in Dulbecco’s Modified Eagle Medium (DMEM) supplemented with 10% fetal bovine serum (FBS), 50 µg/mL gentamicin, and 0.33 µg/mL amphotericin B. Jurkat T cells were maintained in RPMI 1640 medium supplemented with 10% FBS, 50 µg/mL gentamicin, and 0.33 µg/mL amphotericin B.

### Construction of barcoded vectors and virus production

Barcoded vector DNA templates were generated as previously described^17^. Briefly, the GPV^−^ vector was digested with ClaI and MluI, yielding an 11.4-kb DNA fragment lacking the plasmid backbone. A 304-bp barcoded insert was generated by PCR using Phusion High-Fidelity (HF) DNA polymerase (New England BioLabs, Ipswich, MA, USA) according to the manufacturer’s protocol, with primers 51-GACAAGATATCCTTGATCTGNNNNNNNNNNNNNNNNNNNNGCCATCGATGTGGATCTACCACACACAAGGC-31 and 51-CGGTGCCTGATTAATTAAACGCGTGCTCGAGACCTGGAAAAAC-31. The digested GPV^−^ backbone and barcode-containing PCR fragment were assembled using HiFi DNA Assembly Mix (New England BioLabs) at a 1:5 molar ratio.

The resulting circular DNA was cotransfected with pHEF-VSV-G (Addgene #22501) into HEK293T cells using Lipofectamine 2000 (Thermo Fisher Scientific). Plasmid DNA was mixed with Lipofectamine 2000 at a ratio of 1 µg DNA to 2 µL Lipofectamine in Opti-MEM, vortexed briefly, and incubated for 15 min at room temperature before being added to cells in fresh complete medium. Eight hours post-transfection, the medium was replaced with 5 mL of fresh complete medium, and viral supernatants were harvested 48 h later by filtration through a 0.22-µm filter (09-720-511, Fisher Scientific). Viral titers were determined by colony formation assay in HEK293T cells. Following puromycin selection, puromycin-resistant colonies were counted and viral concentrations were calculated as colony-forming units per milliliter (CFU/mL).

### Generation of barcoded proviral pools

Jurkat T cells (5 × 10^6^) were infected at a multiplicity of infection (MOI) of 0.0005, as determined by colony formation assay. Briefly, virus-containing supernatant at the appropriate titer was mixed with Polybrene to a final concentration of 0.5 µg/mL and brought to a final volume of 2 mL with complete RPMI 1640 medium before being added to Jurkat cells. Cells were incubated at 37°C with 5% CO_2_ for 24 h. Viral supernatants were then replaced with fresh complete medium, and cells were incubated for an additional 24 h. Puromycin was subsequently added at a final concentration of 0.5 µg/mL and maintained for approximately 8–10 days, until complete cell death was observed in the uninfected control. Following puromycin selection, infected cells were cultured and expanded until analysis by flow cytometry was initiated.

### Flow Cytometry and Cell Sorting

For flow cytometric analysis, infected Jurkat T cells were pelleted and resuspended in phosphate-buffered saline (PBS) supplemented with 2% fetal bovine serum (FBS) (FACS buffer). Data was collected using a BD LSRFortessa flow cytometer (BD Biosciences) with GFP detected in the fluorescein isothiocyanate (FITC) channel. Flow cytometry data were analyzed using FlowJo single-cell analysis software (version 10.6; BD Biosciences, San Jose, CA). Barcoded proviral pools were sorted by fluorescence-activated cell sorting (FACS). Jurkat T cells were resuspended in FACS buffer and sorted into GFP-positive (GFP^+^) and GFP-negative (GFP^−^) subpopulations using a Invitrogen Bigfoot Cell Sorter (BD Biosciences, Franklin Lakes, NJ). Immediately following sorting, cells were pelleted, resuspended in fresh complete RPMI medium, and expanded until sufficient cell numbers were obtained for downstream analyses.

### Barcode sequencing libraries

Barcodes were amplified from the genomic DNA of infected cells. Genomic DNA was isolated from an aliquot of 2 × 10^6^ infected cells using the DNeasy Blood and Tissue Kit (Qiagen, Germantown, MD) according to the manufacturer’s instructions. Barcode regions were amplified by PCR from multiple reactions containing 100 ng of genomic DNA template using Phusion Hot Start DNA polymerase (New England BioLabs). Primers 5⍰-ACACTCTTTCCCTACACGACGCTCTTCCGATCTGCCTGGCTAGAAGCACAAGA-3⍰ and 5⍰-GACTGGAGTTCAGACGTGTGCTCTTCCGATCTTGCCAATCAGGGAAGTAGCC-3⍰ were designed to flank the barcode region and incorporate Azenta Amplicon EZ adapter sequences. PCR reactions were cycled for 29 cycles with an annealing temperature of 59°C and a 15-second extension at 72°C. Amplicons derived from the same sample were pooled and purified using the Monarch PCR Cleanup Kit (New England BioLabs) prior to submission for high-throughput sequencing using the Azenta Amplicon EZ platform.

### Barcode quantification and analysis

Barcodes were identified and quantified from Azenta Amplicon EZ sequencing data using a previously described custom suite of tools implemented in Python (https://github.com/KiddLab/hiv-zipcode-tools). Briefly, 2 × 75-bp paired-end reads were merged using FLASH v1.2.11. Barcodes were identified by searching for known flanking sequences, and only candidate barcodes with lengths between 17 and 23 nucleotides were retained for analysis. Read counts for each barcode were tabulated, and related barcodes were clustered into families based on sequence similarity. Abundances were then determined by assigning each unique barcode to its most abundant family. The percentage of GFP-positive cells associated with each barcode was calculated as: Fi = [(Gi × P) / (Gi × P + Wi × Q)] × 100, where Fi represents the percentage of GFP^+^ cells for barcode *i*, Gi is the fractional abundance of barcode *i* in the GFP^+^-sorted pool, Wi is the fractional abundance of barcode *i* in the GFP^−^-sorted pool, P is the fraction of total cells sorted into the GFP^+^ pool, and Q is the fraction of total cells sorted into the GFP^−^ pool. Pool reconstruction was done following a second sort of the initial sorted subpools and weighting the calculated GFP+ and GFP− subpool values to the initial unfractionated pool.

### Statistics

Data management and downstream analyses were performed using Jupyter Notebook and graphs were generated using Prism version 10.4.1 (GraphPad). Statistics between timepoints were performed using Mann-Whitney U tests. Associations between replicate pools or between reconstructed and unsorted populations were assessed using Spearman rank correlation coefficients (ρ). Statistical significance was defined as p < 0.05. Exact p values, sample sizes, and tests used are reported in the figure legends.

## Acknowledgements

This study was supported by the Molecular Mechanisms in Microbial Pathogenesis Training Program (T32 AI007528) and the Center for Structural Biology of HIV RNA (CRNA) (U54 AI70660-01).

